# Evaluating the Radiation Sensitivity Index and 12-chemokine gene expression signature for clinical use in a CLIA laboratory

**DOI:** 10.1101/2024.09.19.613957

**Authors:** Anders Berglund, John Puskas, Sean Yoder, Andrew T. Smith, Douglas C. Marchion, Dahui Qian, James J. Mulé, Javier F. Torres-Roca, Steven A. Eschrich

## Abstract

**Background:** The radiation sensitivity index (RSI) and 12-chemokine gene expression signature (12CK GES) are two gene expression signatures (GES) that were previously developed to predict tumor radiation sensitivity or identify the presence of tertiary lymphoid structures in tumors, respectively. To advance the use of these GES into clinical trial evaluation, their assays must be assessed within the context of the Clinical Laboratory Improvement Amendments (CLIA) process.

**Methods:** Using HG-U133Plus 2.0 arrays, we first established CLIA laboratory proficiency. Then the accuracy (limit of detection and macrodissection impact), precision (variability by time and operator), sample type (surgery vs. biopsy), and concordance with reference laboratory were evaluated.

**Results:** RSI and 12CK GES were reproducible (RSI: 0.01 mean difference, 12CK GES 0.17 mean difference) and precise with respect to time and operator. Taken together, the reproducibility analysis of the scores indicated a median RSI difference of 0.06 (6.47% of range) across samples and a median 12CK GES difference of 0.92 (12.29% of range). Experiments indicated that the lower limit of input RNA is 5 ng. Reproducibility with a second CLIA laboratory demonstrated reliability with the median RSI score difference of 0.065 (6% of full range) and 12CK GES difference of 0.93 (12 % of observed range).

**Conclusions:** Overall, under CLIA, RSI and 12CK GES were demonstrated by the Moffitt Cancer Center Advanced Diagnostic Laboratory to be reproducible GES for clinical usage.

## 1 Background

Molecular gene expression signatures (GES) have been developed for multiple purposes within cancer research and for potential use in clinical oncology indications across different tumor types, such as breast [1] and prostate [2] cancers. Many studies use clinical specimens for identifying molecular differences with respect to patient outcome or treatment response. With the advent of several clinically validated gene expression array platforms, new GES can be derived from data generated from these platforms. Far fewer publications exist for evaluating specific GES for clinical decision making and/or clinical trial designs in a Clinical Laboratory Improvement Amendments (CLIA) laboratory setting. The moving of a molecular GES from research-grade data to a reproducible, clinical GES involves experimental evaluation of multiple analytical variables, including sample type and RNA quality [3]. Technical variability is common in molecular GES research studies [4]. However, this variability can be explicitly evaluated in the context of a clinical laboratory that generates the necessary operating characteristics for molecular gene expression tests to be used under clinical conditions. Without adequate assessment of technical variability, clinical validation as a necessary step within the context of a clinical trial, would not be expected to be successful.

The radiation sensitivity index (RSI) is a molecular signature to predict tumor sensitivity to radiation therapy. RSI was developed from a subset of the NCI-60 cell lines that were profiled using Affymetrix HU6800 gene expression arrays [5]. Survival fraction at 2 Gy (SF2) was used with baseline gene expression levels to model an association of gene expression and radiation response. Ten hub genes from among the significant gene expression results were identified using a network biology approach [6]. These 10 genes were combined into a simple rank-based linear model for predicting radiation sensitivity (RSI). This model was demonstrated to predict clinical outcome in patient cohorts [6]. Subsequently, RSI has been used to assess clinical outcome in several cancer types and its radiation-specific nature is indicated in cohorts in which RSI is prognostic only in RT-treated patients [7–16]. Importantly, the RSI model originally developed in 2009 has not been modified from the original formula. RSI was used as the basis for GARD [17], or the Genomically Adjusted Radiation Dose, to estimate the effect of radiation on a tumor by using the linear-quadratic model. GARD has been demonstrated to predict for radiation response more accurately than RSI alone in several diseases, including breast cancer and lung cancer [18–20]. Table 1 highlights the major studies that demonstrated the clinical utility of RSI in retrospective clinical cohorts.

**Table 1.** Publications on the use of RSI/12-CK in retrospective cohorts.

|  | Disease | Reference |
| --- | --- | --- |
| RSI | Breast Cancer | Eschrich et al [7], Torres-Roca et al[10], Sjöström et al.[12], Kang et al. [15] |
|  | Prostate Cancer | Thiruthaneeswaran et al[23] |
|  | Colon Cancer | Ahmed et al[24] |
|  | Sarcoma | Yang, et al[25] |
|  | Penile Cancer | Johnstone, et al [26] |
|  | Pancreatic Cancer | Strom, et al[27] |
|  | GBM | Ahmed,et al[9] |
|  | Lung Cancer | Scott, et al[20] |
|  | Endometrial Cancer | Mohammadim et al[14] |
|  | Bladder Cancer | Khan, et al[28] |
|  | Melanoma | Strom, et al[11] |
|  | Pancancer | Scott, et al[19], Ahmed, et al[29], Strom, et al[27], Ahmed, et al[30], Scott, et al[17] |
| 12CK<br>GES | Colorectal carcinoma | Coppola et al.[21] |
|  | Pan-Cancer/Melanoma | Messina et al.[22] |
|  | Breast | Prabhakaran et al.[31] |
|  | Bladder | Li et al. [32] |
|  | TCGA | Li et al. [33] |
|  | Clear cell renal cell carcinoma | Xu et al. [34] |

The 12 chemokine gene expression signature (12CK GES) (*CCL2, CCL3, CCL4, CCL5*, CCL8, *CCL18, CCL19, CCL21, CXCL9, CXCL10, CXCL11*, and *CXCL13*) was first developed in a large cohort of colorectal cancer samples [21] and showed a strong correlation between the 12CK GES and the presence ectopic lymph node-like/tertiary lymphoid structures (TLS) [22]. The 12CK GES was derived from a set of related chemokines that demonstrated its predictive ability for immunotherapy response, better patient survival, and the presence of TLS, which has been validated in at least 6 cohorts (Table 1).

Given the literature support for the value of these two GES, there is translational oncology interest in evaluating them in prospective clinical trials to determine their clinical utility. To do so, the operating characteristics of the RSI/12CK GES, above and beyond the platform characteristics, must be established. Therefore, we undertook a comprehensive series of experiments to establish these operating characteristics and validation in a Clinical Laboratory Improvement Amendment (CLIA)-certified laboratory (i.e. the Moffitt Advanced Diagnostics Laboratory).

## 2 Methods

Tissue samples were assayed in various conditions using the HG-U133Plus 2.0 GeneChip between 04/02/2019 and 8/17/2020. The Affymetrix GeneChip Scanner 3000 7G was used for MCC experiments. Unless indicated in the experiment, the evaluated samples were generated from 100ng input RNA and 15ug cRNA. The ThermoFisher GeneChip 3’ IVT PLUS reagent kits were used for sample processing.

The most relevant experiments are presented as results and detailed in Table 2. Note that the same GeneChip may be reused in different experiments (e.g., variation over time and operator-to-operator) as appropriate. Supplemental Table 1 includes the specific samples used for each experiment with the corresponding GES (RSI and 12CK GES) values. Tissue/RNA assessment values are included in Supplemental Table 2.

**Table 2.** Experimental design of CLIA validation experiments. The experiments were organized into three categories: Accuracy, Precision and Sample Content. Each of these categories assessed different sources of variability expected to impact signature scores.

| Study | N | Samples | Category | Source<sup>1</sup> | Goal |
| --- | --- | --- | --- | --- | --- |
| **Proficiency** | 8 | 4 samples x 2 labs | Accuracy | CRC | Determine proficiency of CLIA lab by comparing samples from research lab vs. CLIA lab. |
| **Repeatability** | 16 | 4 samples x 4 replicates | Precision | CRC | Determine repeatability of the signature using 4 replicates each of 4 samples run on the same day from the same operator. |
| **Operator** | 12 | 3 samples run in duplicate by 2 operators | Precision | CRC | Determine impact of two different operators |
| **Time** | 8 | 4 samples run in duplicate one week apart | Precision | CRC | Determine impact of processing replicates one week apart |
| **LOD** | 17 | 3 samples were assessed at 5 different amounts of input RNA: 100ng, 25ng, 10ng, 5ng, and 2.5ng.<sup>2</sup> | Accuracy | CRC | Determine a lower threshold of input RNA for successful signature generation. |
| **Macrodissection** | 9 |  | Accuracy | HNC |  |
| **Surgery vs. Biopsy (BRCA)** | 8 | 4 samples x 2 sample types | Sample Content | BRCA | Assess the variability due to sample preparation |
| **Surgery vs. Biopsy (HNC)** | 10 | 5 samples x 2 sample types | Sample Content | HNC | Assess the variability due to sample preparation |
| **MCC vs. External Lab** | 28 | 30 samples attempted x 2 labs<sup>3</sup> | Concordance | HNC | Assess variability from multiple CLIA laboratories |

<sup>1</sup>CRC = colorectal cancer, BRCA = breast cancer, HNC = head and neck cancer
<sup>2</sup>S19/2.5ng was lost when prepared for hybridization.
<sup>3</sup>7 samples failed processing steps; 9 excluded due to poor QC (scaling factor > 1 and percent present < 40%)

### Proficiency Study

Four samples previously profiled in a research genomics facility were profiled in the MCC CLIA laboratory.

### Surgery vs Biopsy Study

We evaluated four breast cancer samples (BRCA) and five head and neck cancer samples (HNC). Core biopsies were obtained from the corresponding tissue block to simulate a clinical biopsy.

### Concordance Study

Thirty frozen tumor specimens (head and neck cancer surgical samples) were profiled in the MCC CLIA Laboratory and an external provider (COV, CLIA Outside Validation). RNA was extracted via the flowthrough for DNA extraction on the Qiacube using the Qiagen AllPrep DNA/RNA/miRNA Universal kit. RNA integrity was assessed using the Tape Station. Due to tissue and array quality concerns, the following filtering criteria were used: MAS5.0 scaling factor *≤* 1, percent present *≥*40 and RIN *≥*6.5. See Supplemental Table 2 for details on RIN values for all samples.

### The 12CK GES

For the 12CK GES, BrainArray36 HGU133Plus2 Hs ENTREZG.cdf Version 25.0.0 downloaded from http://mbni.org/customcdf/25.0.0/entrezg.download/HGU133Plus2_Hs_ENTREZG_25.0.0.zip on 2022-02-25 was used. The following probesets were used for each of the 12CK genes: *CCL2* (6347 at), *CCL3* (missing), *CCL4* (6351 at), *CCL5* (6352 at), *CCL8* (6355 at), *CCL18* (6362 at), *CCL19* (6363 at), *CCL21* (6366 at), *CXCL9* (4283 at), *CXCL10* (3627 at), *CXCL11* (6373 at), and *CXCL13* (10563 at). A PCA model was derived using the 74 samples from the GSE15605 dataset [35]. The raw CEL files were downloaded and processed using IRON [36]. The HGU133Plus2 Hs ENTREZG 25.0.0 version of CDF was used. IRON was used with default settings and GSM390277.CEL was used as median sample. The 11 probesets were selected and a PCA model was calculated. PC1 explains 64.3% of the variation and the PC1/PC2 ratio is 5.1 indicating a robust PCA model [37]. All loadings in the first component are positive indicating that the PCA model behaves as expected.

The 12CK GES scores and RSI scores can be found in Supplemental Table 1 for each dataset/-experiment. The MATLAB code for generating all the figures is available at https://github.com/aebergl/12CK_RSI_Article.

### RSI

Each experiment was normalized using RMA independently using R/Bioconductor. The ten genes were rank-ordered per sample and RSI calculated as previously described in Equation 1. The following probesets were used for each of the RSI genes: *AR* (211110 s at), *JUN* (201466 s at), *STAT1* (AFFX-HUMISGF3A/M97935 MA at), *PRKCB* (207957 s at), *RELA* (201783 s at), *ABL1* (202123 s at), *SUMO1* (208762 at), *CDK1* (205962 at), *HDAC1* (201209 at), and *IRF1* (202531 at).

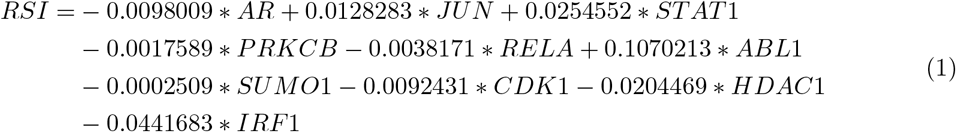

## 3 Results

### Profiling of paired samples in research lab and CLIA lab show high concordance (Proficiency Study)

Four fresh-frozen samples that were previously profiled on the same platform in a research molecular genomics shared resource facility (MGC) were repeated within the Clinical Laboratory Improvement Amendment (CLIA)-certified laboratory. The gene signatures were calculated from arrays independently from the MGC and CLIA environments and the correlation of the signature scores were compared (Figure 1A). Interestingly, the 12CK GES showed a systematic shift in score range but otherwise demonstrated very high correlation (r=0.991). The RSI scores showed less correlation (r=0.762) however the range of observed RSI was compressed in the CLIA experiment. These results emphasize the need to characterize performance of each GES, as the characteristics differ even with the same samples.

**Fig. 1.**
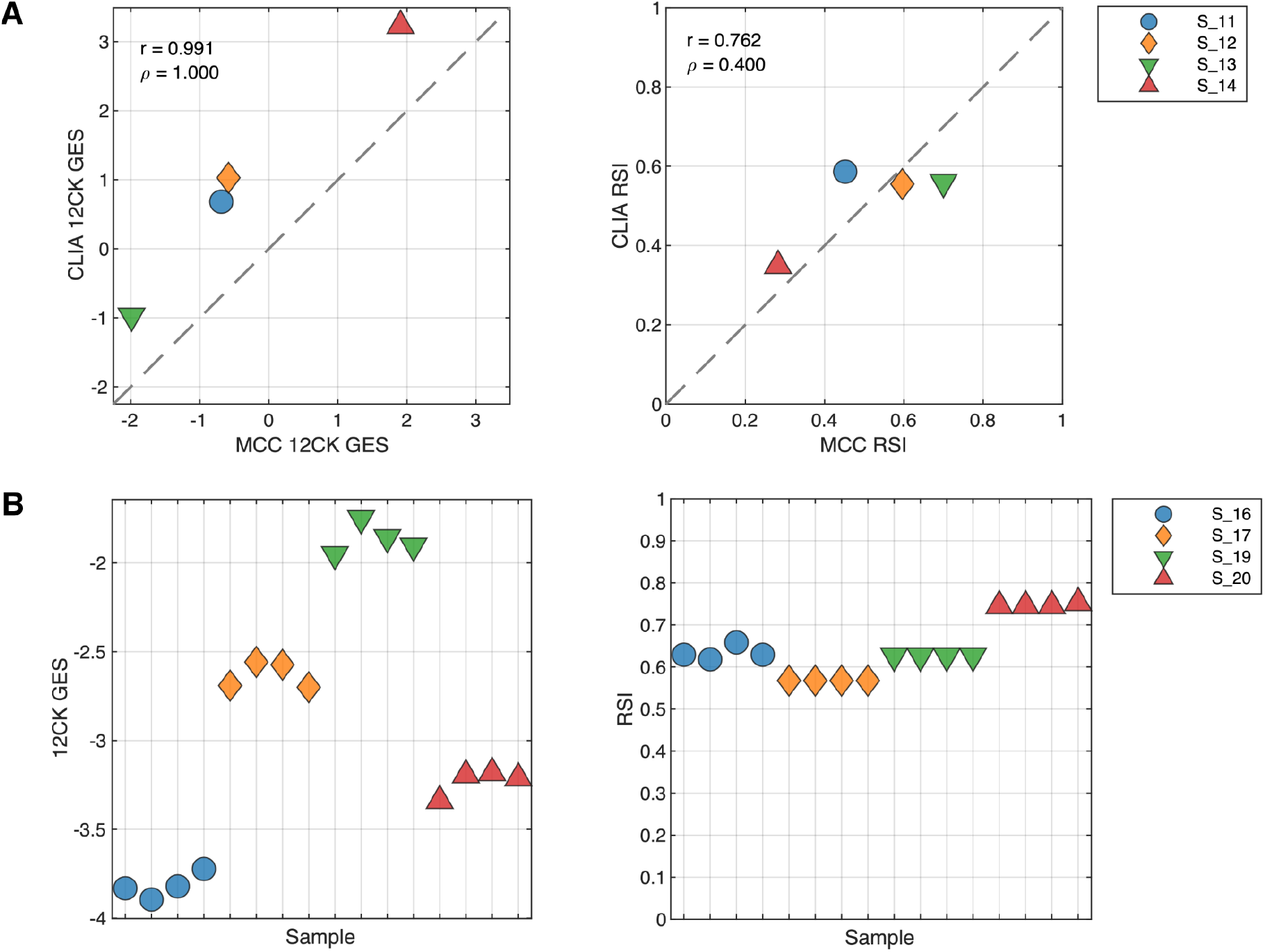
Proficiency and repeatability of the CLIA laboratory in generating 12CK GES and RSI. **(A) Proficiency of CLIA laboratory compared to an established research-grade molecular genomics shared resource facility**. Four samples were previously processed by the MCC Molecular Genomics Core (MGC) and available for CLIA lab processing. GES were derived from both experimental conditions. The experiment was performed to determine that the CLIA laboratory was proficient in generating the expression data for the HG-U133+ platform. (Left) 12CK GES scores in MGC vs. CLIA lab (r=0.991) indicating high correlation although signature calibration was needed. (Right) RSI signature scores in MGC vs. CLIA laboratory indicating compressed RSI signal from the CLIA experiments. **(B) Repeatability of GES from quadruplicate samples in CLIA laboratory**. Four samples were processed in quadruplicate and arrayed in the CLIA laboratory from the same operator. GES scores were derived from each experiment. (Left) 12CK GES scores had low variability in each of the four samples. (Right) RSI was identical in two samples and had low level of variability in two samples.

### Replicated assays demonstrate repeatability of signature scores

We next profiled four samples in quadruplicate to assess the repeatability of the GES. Each sample was processed independently by the same operator resulting in four distinct gene expression arrays per sample. As shown in Figure 1B, the 12CK GES varied for each sample but the overall variability was low with a mean range of scores by sample of 0.16825 (7.8% of total observed range). Likewise, the RSI score variability was generally low with a mean range of scores by sample of 0.01 (6.5% of total observed range). In the case of RSI, two samples (S 19 and S 17) show identical signature scores across all four replicates. The RSI is rank-based, therefore small variations in expression do not always result in differences in GES score. Operators and time demonstrate higher RSI variability, but not 12CK variability

The replicability and precision of the GES was assessed using two different experiments, evaluating the impact of operator and processing date on the GES. To test the operator characteristics, three samples were run in duplicate by two different operators (O1, O2) (Figure 2A). The 12CK GES showed a very small operator effect, much smaller than the sample differences in GES score. In contrast, the operator had a larger impact in one sample (S 11) for RSI. The impact of processing date was assessed by profiling four samples that were independently processed one week apart (Figure 2B). The 12CK GES showed low variation among replicates across time. In the case of RSI, S11 and S12 did show differences (less than 0.1 difference) in replicates while the other two replicates were identical in score.

**Fig. 2.**
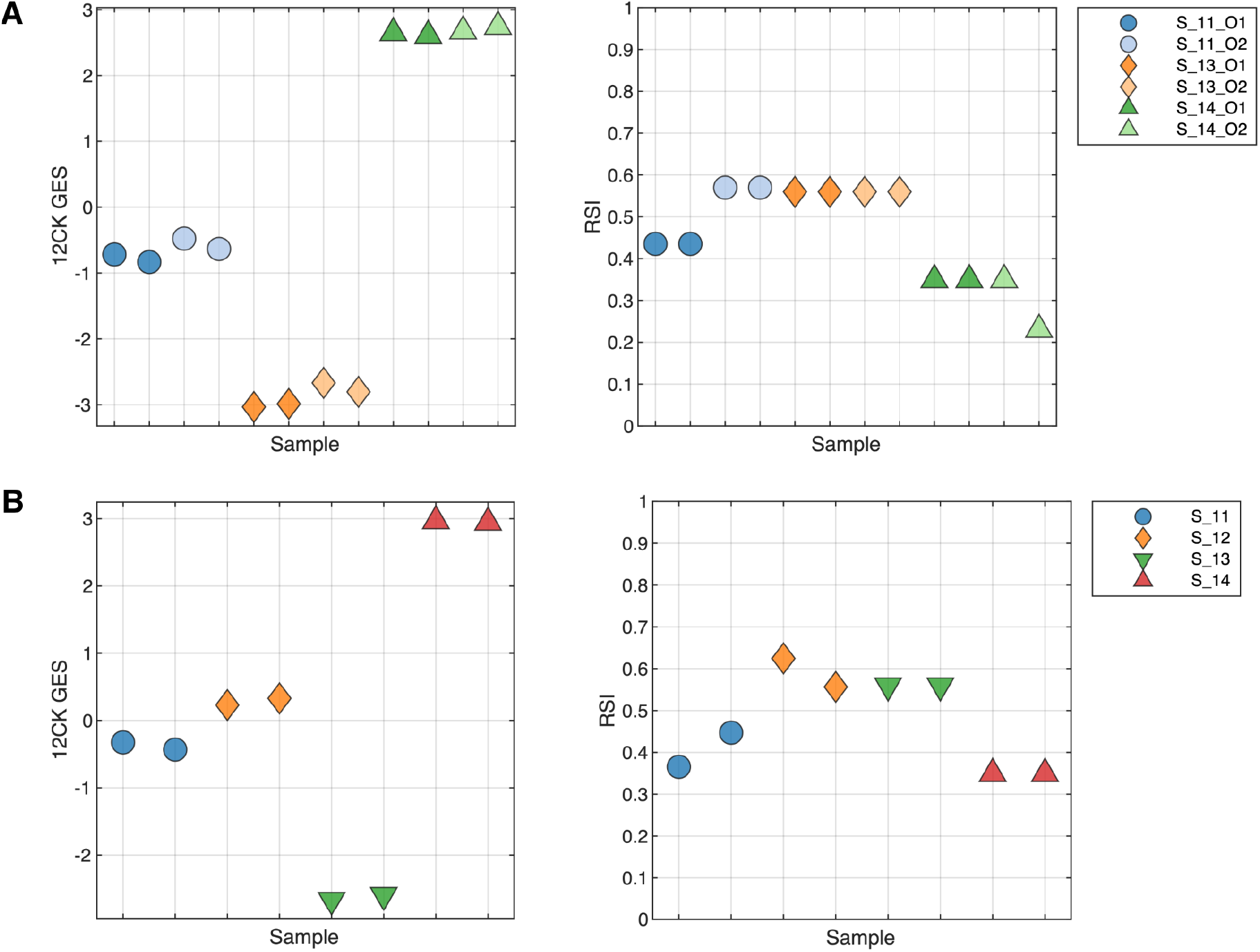
Replicability and Precision Analysis through measuring operator-to-operator and repeated run experiments. **(A) Operator Variability:** Three samples were run in duplicate by two different operators to assess both the variability in operator handling as well as repeatability from the same operator. The 12CK GES showed low variability across operator whereas the RSI score had a larger difference in RSI score between operators. **(B) Repeated run over time:** Four samples were repeated one week apart and assessed. The variability in the 12CK GES was low for all samples. RSI showed variability (less than 0.1) in two of the four samples.

### Summary Reproducibility

Using the experiments in which four replicates were produced (either from operator-to-operator variability or the repeatability study), Table 3 shows the summary mean and standard deviation for the GES scores. Overall, reproducibility analysis of the scores indicated a median RSI difference of 0.06 (6.47% of range) across samples and a median 12CK GES difference of 0.92 (12.29% of range). The GES are very reproducible in the CLIA laboratory using the pre-defined processing protocols.

**Table 3.**
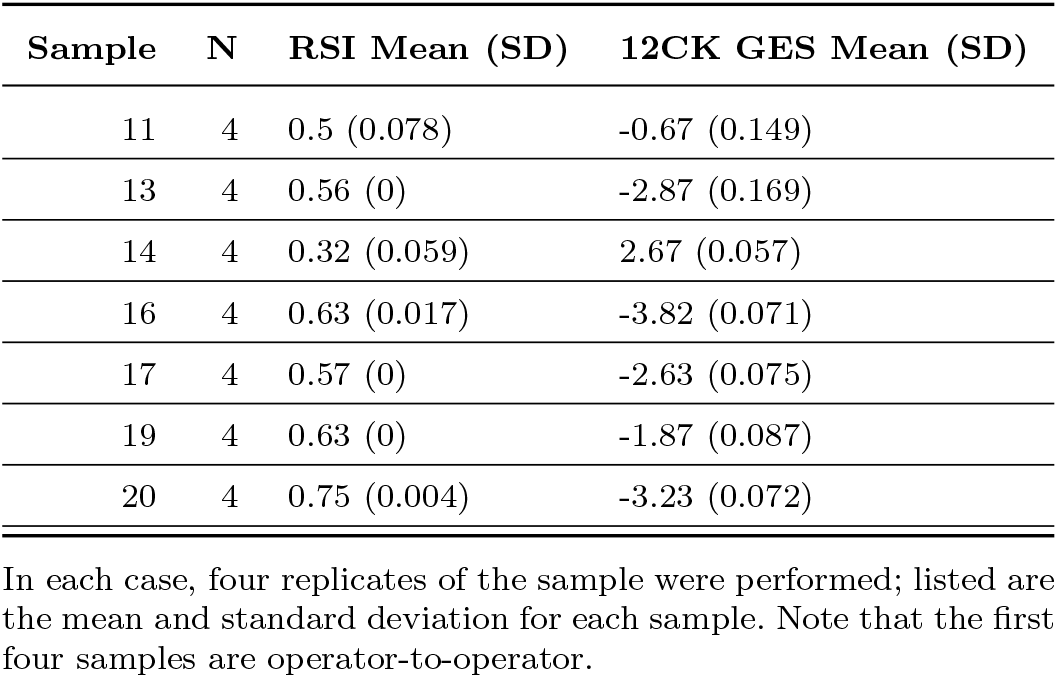
Summary of reproducibility of signature scores across operator to operator and repeatability experiments.

### Amount of material (input RNA) indicates 2.5 ng is lower limit for GES

An important consideration for any molecular test is the amount of material required to perform reliably and robustly. To address this question, three samples were profiled using 100, 25, 10, 5 and 2.5 ng of input RNA (Figure 3). For the 12CK GES, the scores are very similar across RNA amounts, with a lower score observed in the 2.5 ng condition suggesting a lower limit on RNA. RSI demonstrated more overall variability (less than 0.1), however did not appear to have an input amount-related impact.

**Fig. 3.**
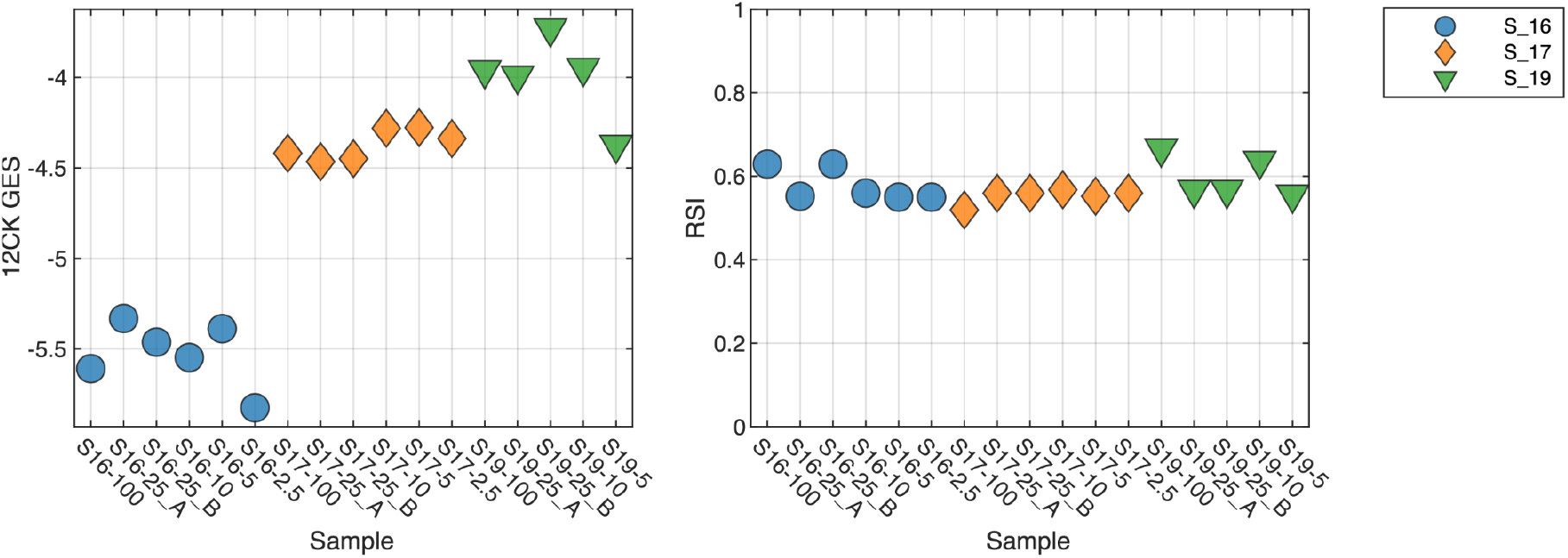
Impact of amount of input RNA on GES scores. Three samples were profiled using differing amount of input RNA (100ng, 25ng, 10ng, 5ng, and 2.5ng). Lower 12CK GES scores were observed at 2.5ng suggesting a lower limit on input RNA. RSI demonstrated more variability overall but did not appear to have a systematic difference at 2.5ng.

### Tissue Type Factors

We also assessed the impact of macrodissection or normal tissue mixtures on GES score. From three samples, we examined the tumor macrodissected tissue (T), normal macrodissected tissue (N) and non-macrodissected tissue (PM). As shown in Figure 4, the normal (N) scores demonstrate large differences from tumor (T) as is the case with the 12CK GES for S1. In this case, the non-macrodissected tissue (PM) 12CK GES score was more similar to the corresponding normal (N). In other cases, the RSI score (S3) showed a large difference between normal and tumor. These results indicate that, as expected, GES scores derived from tumor tissue must be assayed from macrodissected (predominantly) tumor tissue to be reliably reproduced.

**Fig. 4.**
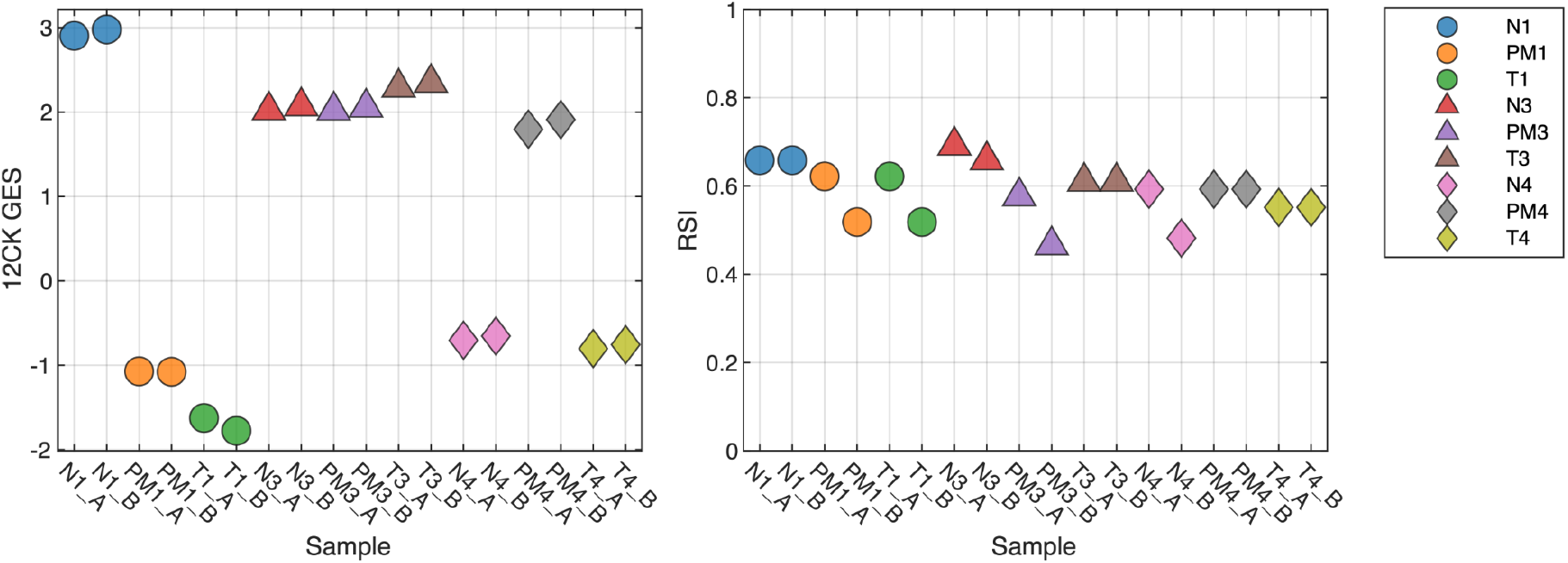
Impact of Macrodissection on signature scores. Three samples were profiled across three different sample conditions: tumor macrodissected tissue (T), normal macrodissected tissue (N) and non-macrodissected tissue (PM). As expected, normal tissue can result in large changes to the signature score, which can be seen in non-macrodissected tissues as well. For instance, the 12CK GES score for Normal tissue from sample 1 is much higher however there is elevated signal in the PM sample as well, whereas the differences in sample 3 (N,T,PM) are small.

### Surgical specimens vs. punch biopsies in breast cancer and head & neck cancer introduces variability

While both GES were developed from macrodissected surgical specimens, punch biopsies are often a more readily available source of material for clinical assays. Therefore, we assessed the variability in GES scores between these preparation types in two different diseases: breast cancer and head & neck cancer. Core biopsies were obtained from tissue blocks. Figure 5A indicates that in breast cancer, the 12CK GES score can be attenuated by punch biopsy whereas the RSI score showed small changes in both directions. In the case of head & neck cancer, both the 12CK GES and RSI scores showed increases in scores for biopsy samples (Figure 5B). This may relate to the cellular content and/or proportion of immune cells present in the specimen. The variability introduced into the scores due to preparation type can be used when considering alternative methods for use. Interestingly, RSI tended to have a much narrower range of values (e.g., 0.6-0.8 for breast cancer vs 0.2-0.5 head & neck cancer) indicating some tissue-specific sensitivity that has been noted previously.

**Fig. 5.**
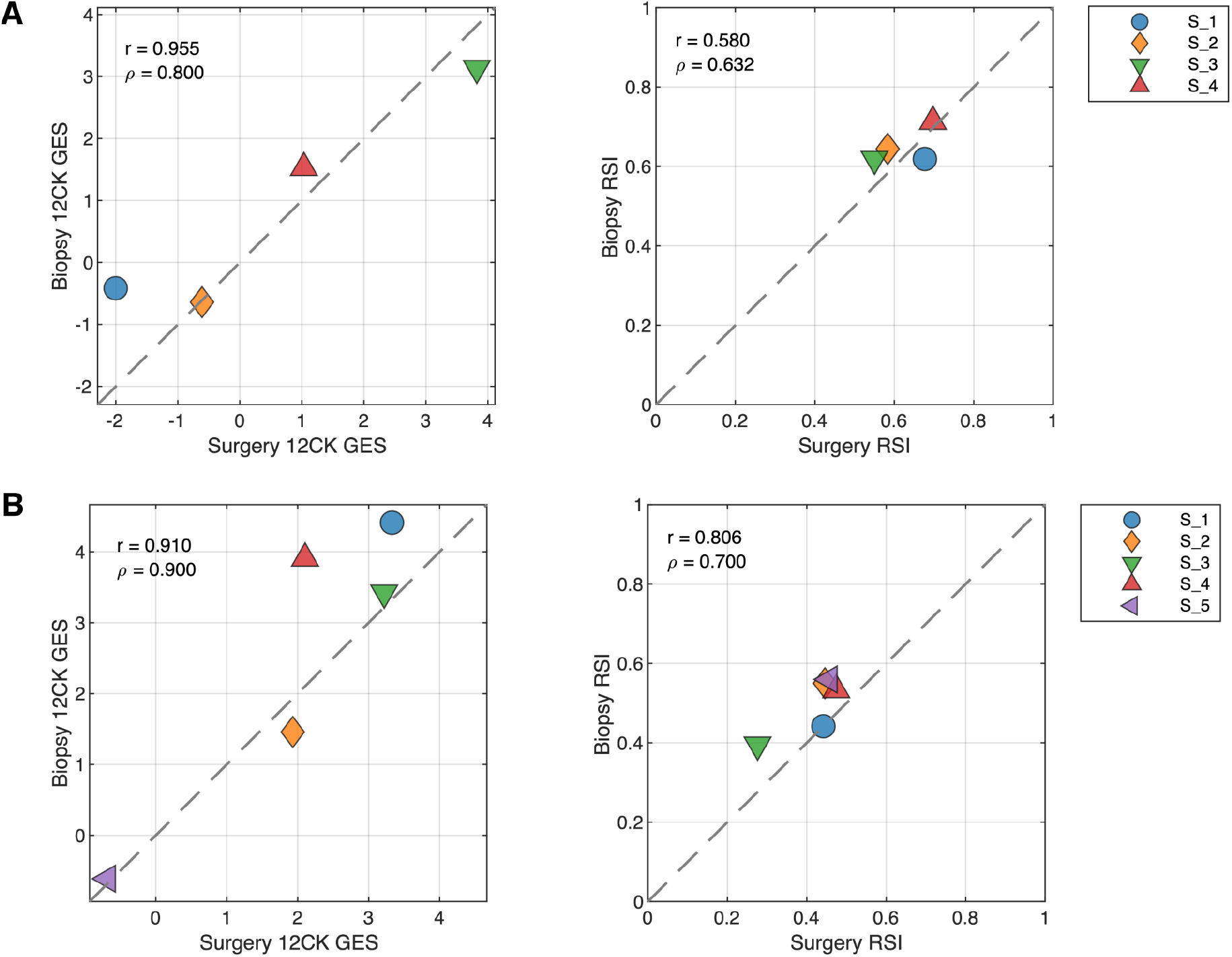
Impact of Surgery vs Biopsy Sample on Signature Scores. (A) Breast cancer specimens from tissue resection (surgery) and punch biopsy (biopsy) were compared. (B) Head and Neck cancer specimens from tissue resection (surgery) and punch biopsy (biopsy) were compared.

### Concordance with External Laboratory

Thirty samples were processed internally at the Moffitt Cancer Center (MCC) CLIA Laboratory and sent to an external vendor for processing. Five samples failed processing and were not hybridized; two samples where hybridized but failed initial QC. An additional 9 samples were excluded due to poor QC. After filtering, GES scores from 14 samples were compared between the MCC CLIA Laboratory and the external vendor (Figure 6). Using the Passing-Bablok test [38], both the 12CK GES and RSI signatures had linear relationships between the two sites. The median RSI score difference was 0.065 (6% of full range) and the 12CK GES difference was 0.93 (12% of observed range).

**Fig. 6.**
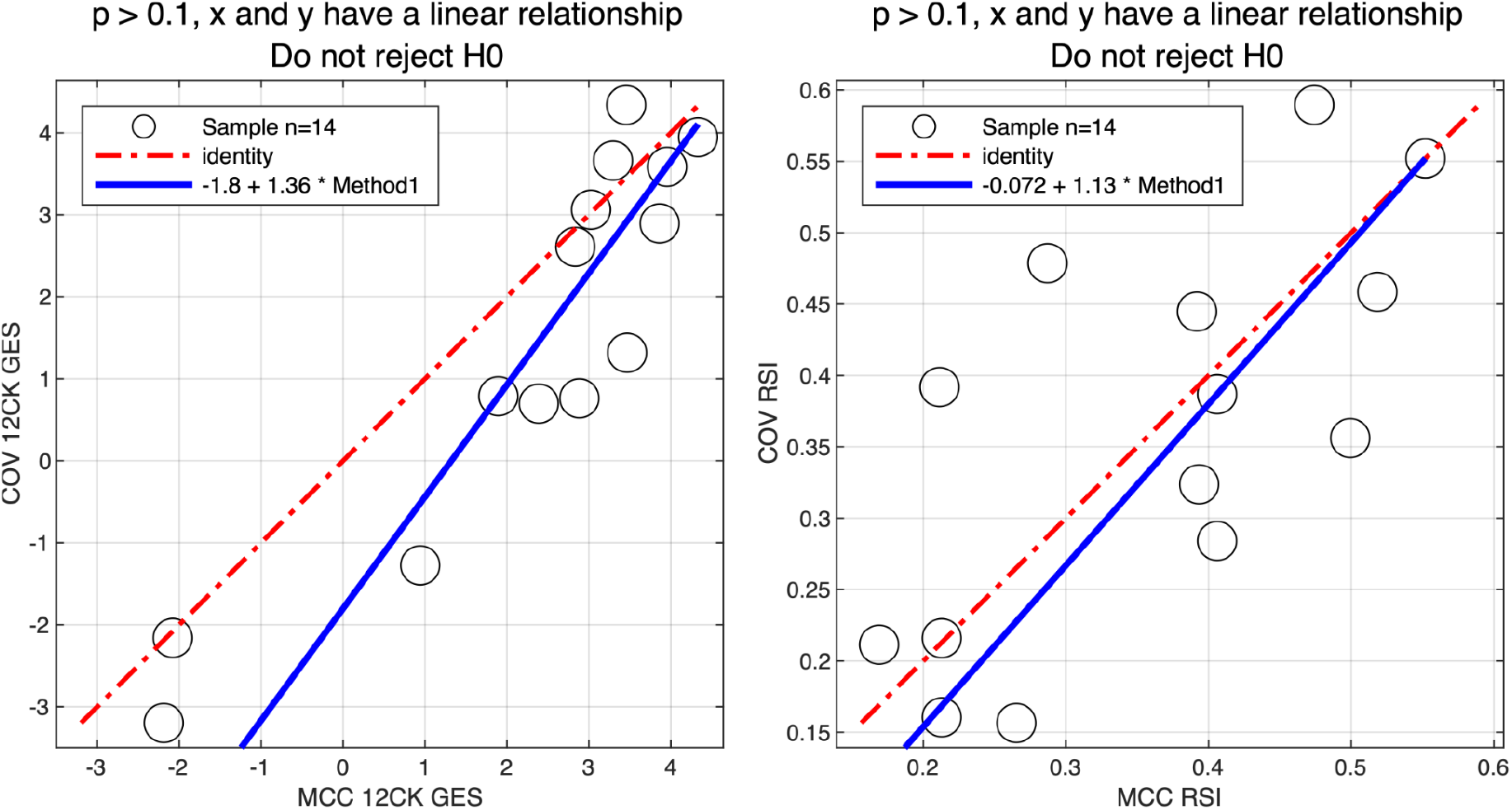
Concordance of signature scores across MCC CLIA Laboratory and external vendor. 30 samples were profiled in the MCC CLIA Laboratory and external vendor. After excluding low quality samples, the linearity of the 12CK GES (left) and RSI (right) scores for 14 samples across sites was assessed.

## 4 Discussion

We describe a series of experiments that were performed to systematically assess the assay platform (the Affymetrix HG-U133Plus 2.0 GeneChip) and specific gene signatures (RSI and 12CK) for CLIA laboratory use in clinical trials assessing utility. Our results indicate that specific GES have different operating characteristics, even using the same platform and hybridizations. Thus, it is important for each GES to undergo systematic evaluation for its own unique precision and robustness limits.

The studies described herein were initially performed to provide support for the CLIA validation of RSI. We note that a “valley of death” exists for GES being deployed clinically. Retrospective analyses across a variety of platforms are more typically observed in research studies, however clinical translation requires explicit experiments to clearly assess the operating characteristics under controlled conditions. Importantly, the arrays generated in this study were subsequently reused to assess the variability the 12CK GES without requiring additional hybridizations. As new GES are developed, this dataset can support rapid verification any additional GES under controlled conditions. These types of experiments systematically generated across clinical specimens as a public resource can provide initial data on the reliability of the GES for clinical translation, in particular where patient decisions are made.

One aspect for GES is the underlying model used for prediction. RSI, for instance, was developed as a rank-based linear regression to maximize robustness across multiple platforms. This approach has been used for validation in many retrospective cohorts. Notwithstanding, the model was not designed to optimize precision. This can be seen in the evaluation for CLIA operations; the level of variability observed for RSI is higher than the 12CK GES for this reason. In contrast, the 12CK GES was developed as a PCA-based model and is likely more robust to individual genes introducing variability. In both cases, the research-grade model has been used for extensive retrospective validation in clinical cohorts. Modifying the scoring algorithm for precision is not possible without re-validating the approach across retrospective datasets. An open area for additional research is a systematic approach for hardening, modifying or adapting an existing model without negating the value of the prior clinical validation data.

## List of Abbreviations

(RSI): Radiation Sensitivity Index
(12CK GES): 12-chemokine gene expression signature
(GES): gene expression signatures
(CLIA): Clinical Laboratory Improvement Amendments
(SF2): Survival fraction at 2 Gy
(GARD): Genomically Adjusted Radiation Dose
(MCC): Moffitt Cancer Center
(COV): CLIA Outside Validation
(TLS): tertiary lymphoid structures
(BRCA): breast cancer
(HNC): head and neck cancer
(CRC): colorectal cancer
(MGC): molecular genomics shared resource facility

## Declarations

### Ethics approval and consent to participate

Informed consent was obtained from patients to the Total Cancer Care Protocol, the Moffitt Cancer Center’s institutional biorepository (MCC#14690; Advarra IRB Pro00014441). Tissue experiments were determined to be non human subject’s research (NHSR) by Moffitt Cancer Center under the Cancer Center’s Biospecimen Pilot Project Process.

### Supplementary information

- **Supplemental Table 1:** RSI/12CK GES scores for all conditions reported.
- **Supplemental Table 2:** Source sample characteristics.

The code for generating the figures and results within the paper are available on github at https://github.com/aebergl/12CK_RSI_Article.

## Authors’ Contributions

JTR designed the study; AB/SAE performed data analysis; JP performed CLIA array experiments; SY and ATS performed MGC experiments; DCM managed tissue banking/acquisition, DQ oversaw CLIA array experiments; JJM/JTR manuscript writing and editing. All authors reviewed the manuscript.

